# Identification of potential natural compound inhibitors and drug like molecules against human METTL3 by docking and molecular dynamics simulation

**DOI:** 10.1101/2022.06.19.496750

**Authors:** Shibambika Manna, Pragati Samal, Rohini Basak, Anushka Mitra, Arijit Kumar Roy, Raima Kundu, Amrita Ahir, Amlan Roychowdhury, Ditipriya Hazra

## Abstract

Nucleotide level chemical modification in transcriptome is critical in regulating different cellular processes, including cancer. The most investigated epitranscriptomic modification is methylation at the N^6^-position of adenosine (m^6^A). This dynamic modification process is carried out by: writer, reader and eraser proteins. Writers are methyltransferases, METTL3 is the major writer that works in association with METTL14, an accessory protein. Extensive study revealed that cancer progression for acute myeloid leukaemia, gastric cancer, colorectal cancer, hepatocellular carcinoma, and lung cancer is directly contributed by irregular expression of METTL3. Targeting METTL3, has opened a new window in the development of new inhibitors/drugs. In this study, 80 commercially available compounds were found from an unbiased screening by molecular docking, showing better score when compared with the existing substrate/substrate-analogue and the inhibitor bound crystal structures in terms of docking score and binding energy calculation. Among this pool of compounds, the best seven small molecules, AMF, RAD, JNJ, MEH, ECP, MHN, SGI, have been selected and further validated by different computational tools like binding energy calculation, molecular dynamics simulation etc. The novel hits found in this study can function as lead compounds which can be developed into inhibitors as well as drugs, specific against METTL3.

## Introduction

DNA and proteins have been researched to undergo various modifications during the course of its lifetime. Similarly in the 1970s, mRNA was also found to be undergoing similar modifications, which have been termed as post-transcriptional modifications. Desrosiers [1] and colleagues discovered m^6^A (N6-methyladenosine) to be the most significant form of modification in mRNA including mammals [2], insects [3], plants [4], yeast [5] and some viruses [6]. Jia et al., [7] found out that m6A modification is ‘dynamic’ and ‘reversible’ with the discovery of the first m6A demethylase - fat mass and obesity-associated protein (FTO). After this discovery, m6A modifications have been reported in several diseases, and bioprocesses [7].

The prospect of m^6^A modifications was significantly established with the sequencing of m^6^A by using Next Generation Sequencing (NGS) [8]. With the sequencing results, it was found that the modifications take place in the consensus sequence - RRACH (in which R represents A or G and H represents A, C or U) and were enriched in the 3’-Untranslated regions along with the stop codons. The mapping studies have revealed very significant functions of m6A transcriptome. Nearly 170 modifications have been reported in RNA including - N6-methyladenosine (m^6^A), N1 -methyladenosine (m^1^A), inosine (I), 5-methylcytosine (m^5^C), and pseudouridine (ψ) [9] - out of which 72 variants of methylation has been recognised which translates to its highly pervasive nature in a transcriptome-wide manner.

Extensive research in this field marked the formal establishment of “epitranscriptomics” that deals with the dynamics and functionality of an mRNA transcriptome [10, 11]. epitranscriptomic modifications have been found to play a crucial role in regulating RNA biology such as splicing, export, maturation, stability, degradation, translation [12,13], folding, and metabolism. In addition to mRNA, other RNAs such as - microRNA (miRNA), long noncoding RNAs (lncRNA), circular RNA (circRNA), ribosomal RNA (rRNA), transfer RNA (tRNA) and small nucleolar RNAs (snoRNA) have also been reported to undergo m^6^A modifications [14,15]. Interestingly, m^6^A methylation does not undergo in cytosolic mRNAs [16,17,18,19]. m^6^A modification has been reported to play a vital role in the maintenance of embryonic stem cell (ESC) pluripotency, regulation of spermatogenesis, regulation of brain development, involvement in the development of nerve cells and neural regulation in adults [20]. Similar links have been found in between the regulation of cancer and epitranscriptomic modifications [21,22,23].

The functions of m^6^A modifications are exercised by ‘writer’, ‘reader’ and ‘eraser’ proteins. RNA demethylases are erasers, RNA methyltransferases are writers, and m6A binding proteins are readers. The first writer protein to be discovered was Methyltransferase 3 also known as METTL3, that weighs about 70 kDa, constitutes of 580 amino acids and its enzymatic activity is executed by zinc finger domain (ZFD) and a methyltransferase domain [24]. It has been found out that METTL14 (another writer protein) plays the structural role for RNA-binding whereas, METTL3 plays the catalytic role.

METTL3 enabled to discover the relationship between m^6^A and cellular physiology [25]. METTL3 was found to be acting as an essential oncogene in Adult Acute Myeloid Leukemia (AML), where METTL3 is recruited to induce m^6^A deposition which in turn [26] plays a crucial role in leukaemia progression by actively engaging translation of target mRNAs [27] and knock-down of METTL3 down-regulates the progression of cancer. Henceforth, METTL3 have been reported to have direct relationship with its overexpression and oncogenic implications in many other types of cancer [28]. In hepatocellular carcinoma (HCC), METTL3 was observed to be significantly upregulated and contributed heavily in the progression of tumorigenicity whereas its depletion resulted in the reduced metastasis [29]. As in case of gastric cancer it was observed that the upregulation of METTL3 promoted cell proliferation and migration. It was directly related to the tumour survival and metastasis [30]. Lung cancer, which is reported to be the most widespread affecting form of cancer, has also been found to have its progression affected by the elevated levels of METTL3. METTL3 induces drug resistance and progression of NSCLC cells by increasing the levels of m^6^A modifications [31]. Li et al. suggested that in case of colorectal cancer (CRC), METTL3 expression is associated with the progression of CRC metastatic tissues [32]. A positive feedback-loop has been found to exist between METTL3 and the expression of HBXIP in breast cancer cells suggesting that METTL3 indeed drives the occurrence of breast cancer [33]. METTL3 have been found to induce hepatoblastoma [34], non-small cell lung cancer [35], bladder cancer [36] and more. Hence, METTL3 is a very lucrative target for cancer inhibition. Finding inhibitors that specifically target METTL3 is an active field of research [37,38]. So far there is only one commercially available inhibitor specific for METTL3, STM2457 [39]. It is essential to find more potent inhibitors and druglike molecules which can be developed into effective drugs in future for cancer treatment.

In this work we have targeted the S-Adenosyl Methionine (SAM)/ S-Adenosyl Homocysteine (SAH) binding pocket for unbiased virtual screening with 810 commercially available small molecules and have found novel molecules, potential inhibitors of METTL3 using molecular dynamics simulation, protein-ligand binding energy calculation and pharmacokinetic analysis.

## Results

### Molecular docking

The docking protocol has been validated by comparing the METTL3-SAH co-crystal (PDB ID: 5IL2, chain A) with docked SAH. The RMSD between the docked and co-crystallized structure was 0.065Å, indicating the orientation of small molecules is very similar in both cases (supplementary Figure 1). The binding energy of docked SAH was −8.7kcal/mol. Initial *in-silico* screening was performed using known DNA methyltransferase inhibitors followed by unbiased screening of commercially available natural products to find potential inhibitors and avoid the prejudice of searching among compounds with similar structure. The SDF files of small molecules were downloaded from PubChem and converted to PDB format using Open Babel. Autodock Vina [40] produces eight docked conformations, among them the one with highest score was selected for further processing. Over 800 molecules were screened. Since the binding energy of the docked SAH was −8.7 kcal/mol, the cut off was set at <-9.0 kcal/mol, to find molecules that have significantly stronger binding. 80 compounds have been found with docking energy equal to or better than −9.0 kcal/mol (supplementary Table 1). The best 10 small molecule inhibitors based on the docking score are tabulated in Table 1 along with a three-letter code, used in this manuscript as a unique identifier, and chemical structure. Fig 1 and Table 2 describes the specific interactions between METTL3 and top 7 inhibitor molecules. The surface representation of METTL3 with the inhibitor molecules clearly exhibits how different inhibitors has occupied the binding pocket and interacting with the target protein. The detailed amino acid level information of the protein-inhibitor interaction is presented in the right panel of Fig 1 and in Table 2. In the table, the nature of interaction (H bond or Hydrophobic interaction) has been tabulated in the amino acid level and the figure is representing the spatial distribution of interacting amino acids in the docked structures (The LIGPLOT analysis for each inhibitor-protein complex is presented in supplementary Figure 2).

**Table 1:**
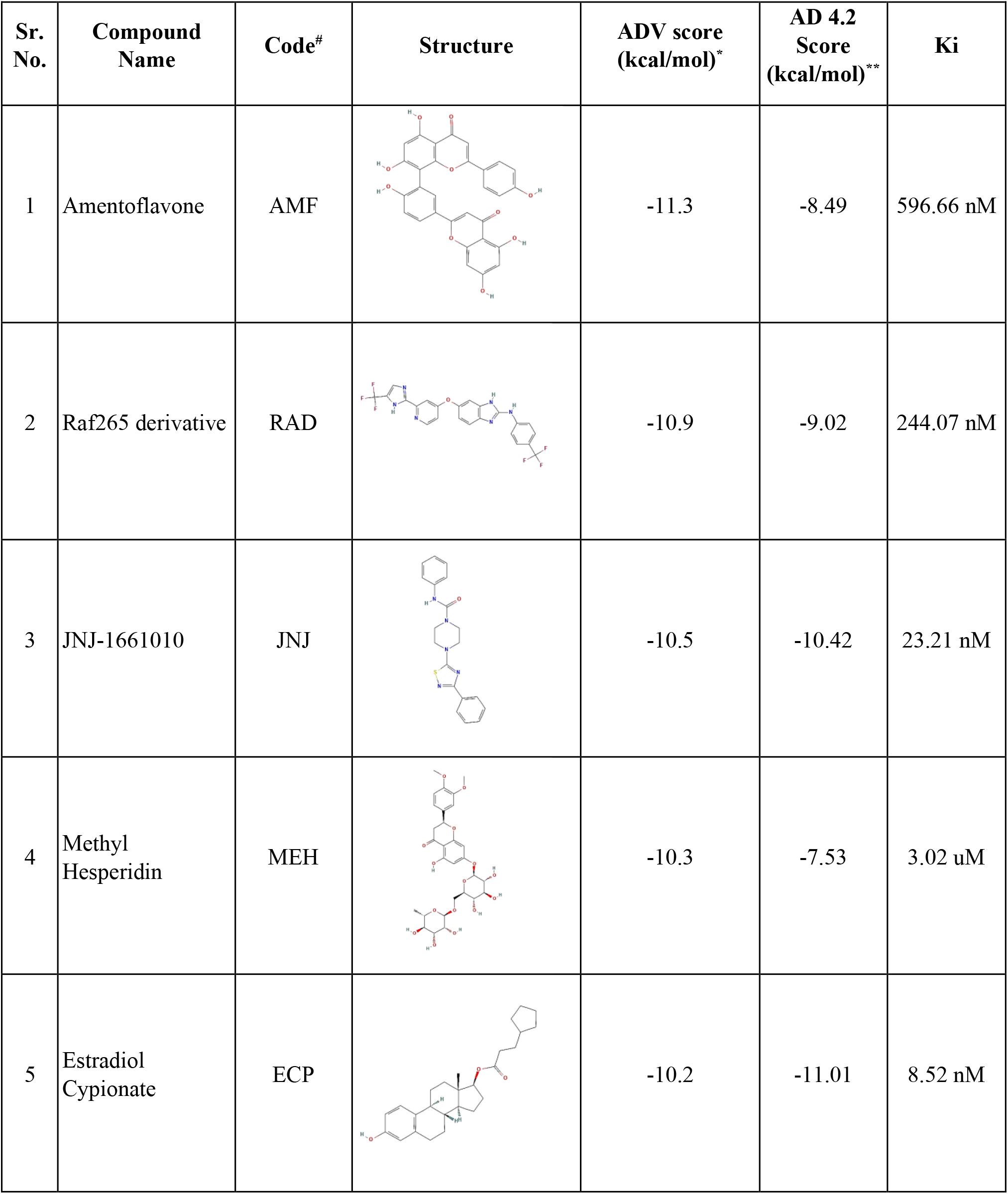
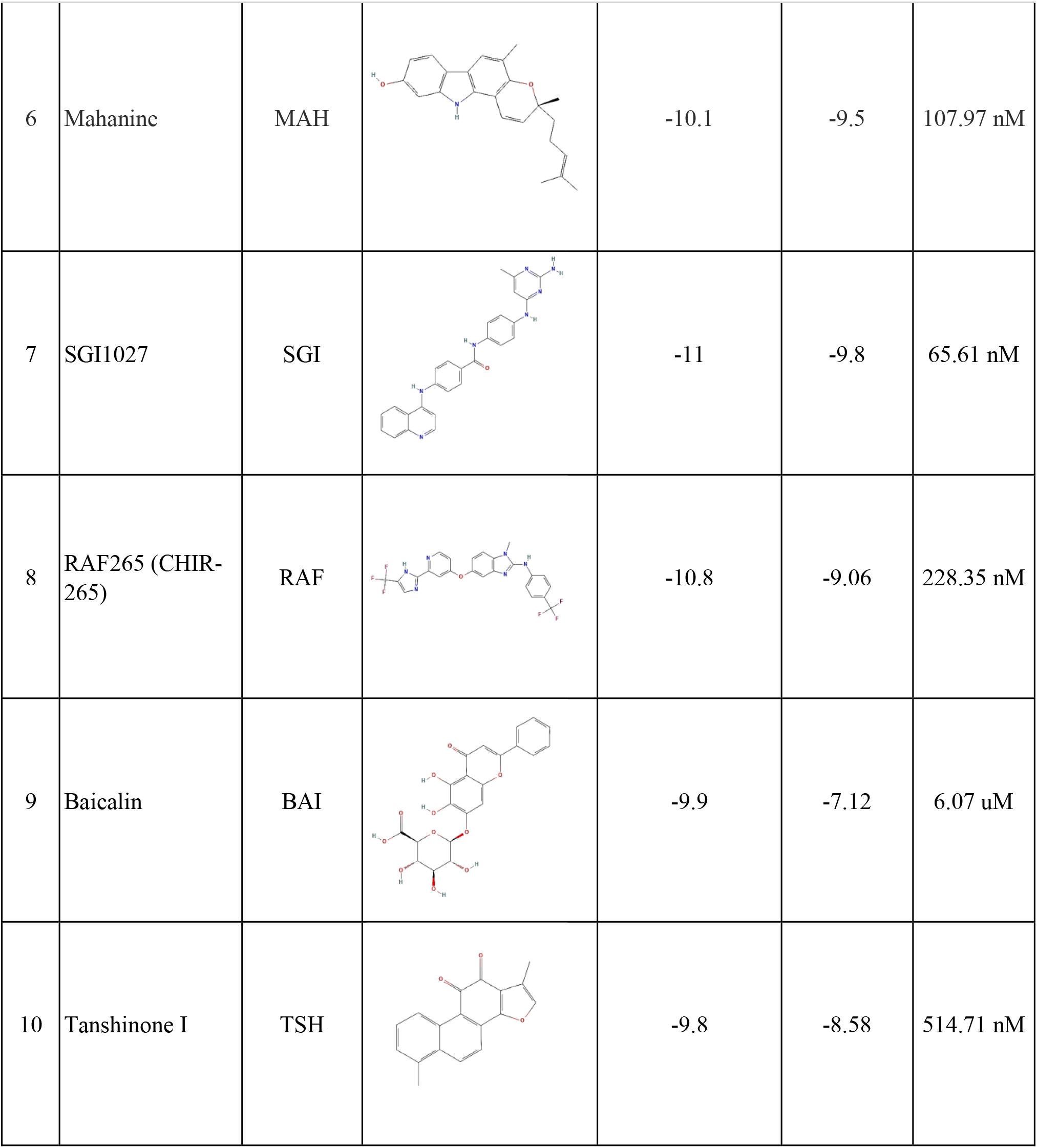
chemical structures and docking scores of the best hits. Details of the best 10 hits with chemical structure, three letter code, docking score (# unique three letter code for the small molecules, *ADV-Autodock Vina and **AD 4.2-Autodock 4.2) and inhibition constant (Ki).

| Sr. No. | Compound Name | Code<sup>#</sup> | Structure | ADV score (kcal/mol)<sup>*</sup> | AD 4.2 Score (kcal/mol)<sup>**</sup> | Ki |
| --- | --- | --- | --- | --- | --- | --- |
| 1 | Amentoflavone | AMF |  | -11.3 | -8.49 | 596.66 nM |
| 2 | Raf265 derivative | RAD |  | -10.9 | -9.02 | 244.07 nM |
| 3 | JNJ-1661010 | JNJ |  | -10.5 | -10.42 | 23.21 nM |
| 4 | Methyl Hesperidin | MEH |  | -10.3 | -7.53 | 3.02 uM |
| 5 | Estradiol Cypionate | ECP |  | -10.2 | -11.01 | 8.52 nM |
| 6 | Mahanine | MAH |  | -10.1 | -9.5 | 107.97 nM |
| 7 | SGI1027 | SGI |  | -11 | -9.8 | 65.61 nM |
| 8 | RAF265 (CHIR-265) | RAF |  | -10.8 | -9.06 | 228.35 nM |
| 9 | Baicalin | BAI |  | -9.9 | -7.12 | 6.07 uM |
| 10 | Tanshinone I | TSH |  | -9.8 | -8.58 | 514.71 nM |

**Figure 1:**
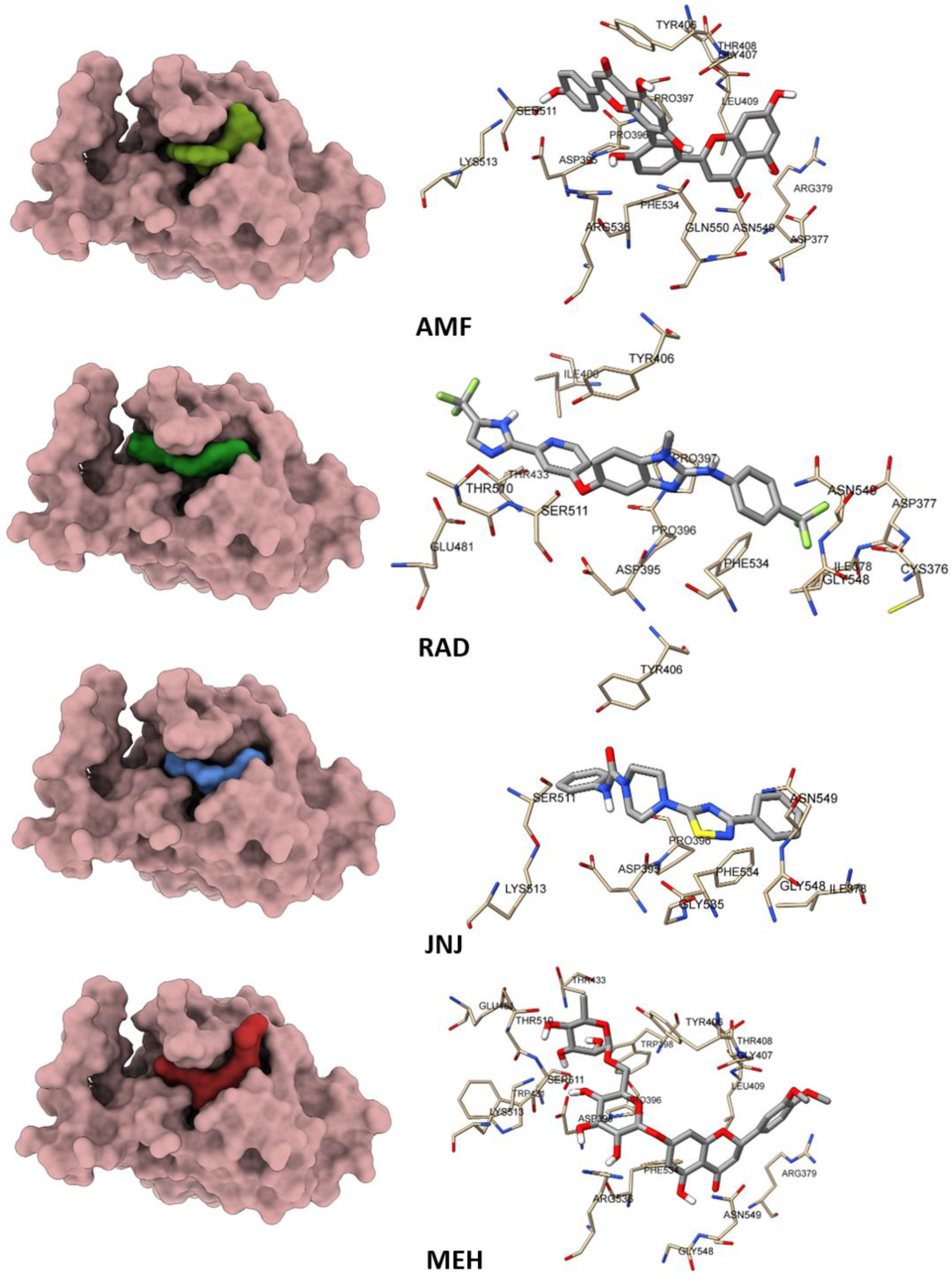
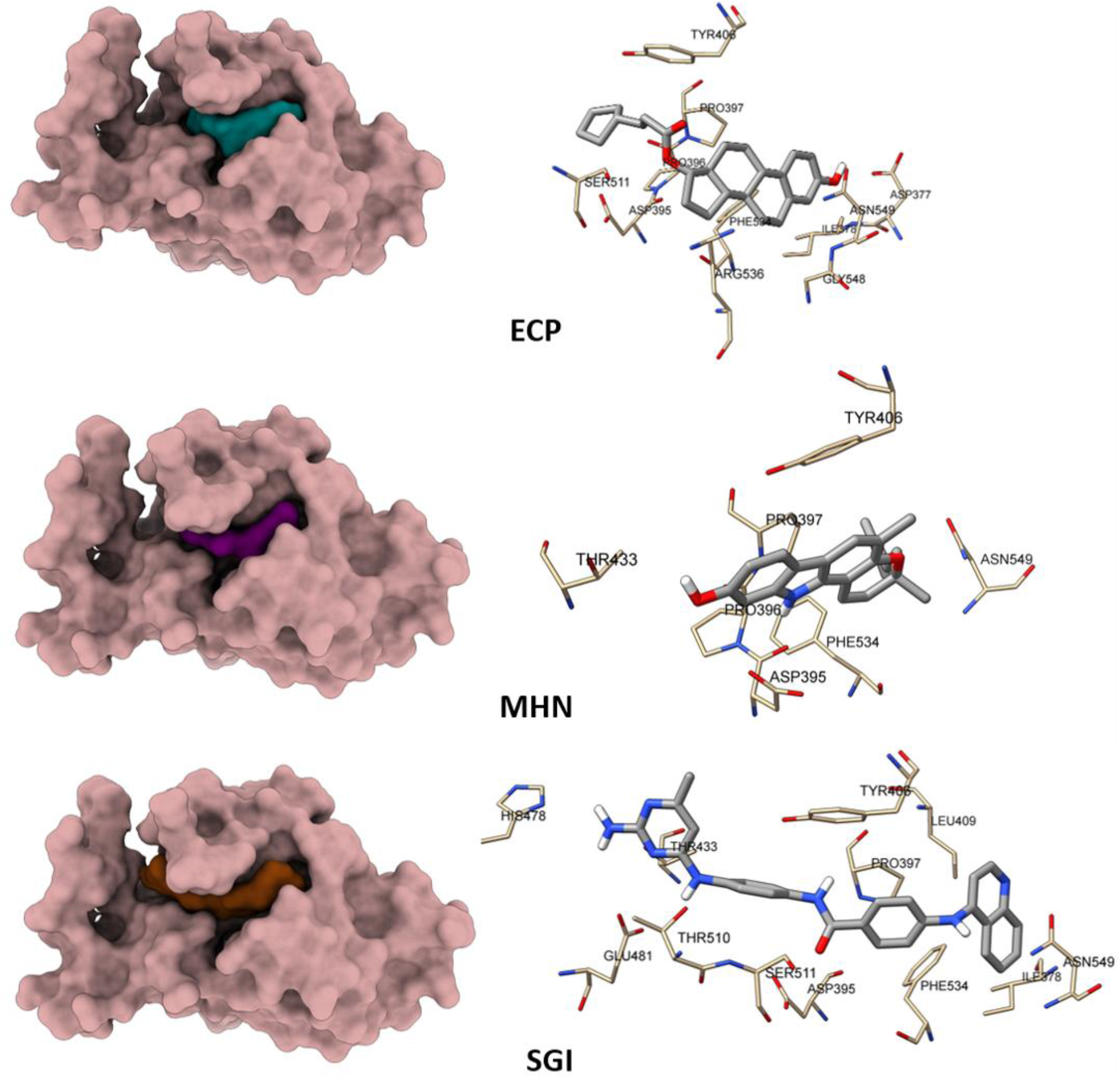
The interaction of small molecule inhibitors with binding pocket of METTL3. The left panel is surface representation of human Mettl3 protein (PDB ID 5IL2) with different small molecule inhibitor bound form which depicts the relative position of the inhibitor molecules in the binding pocket. In the right panel the amino acid residues of the binding pocket which has established interaction with the small molecule inhibitor via either hydrogen bond formation or hydrophobic interaction has been shown in stick mode of representation.

**Table 2:** Interaction between METTL3 and inhibitors. Amino acid residue wise interaction between Mettl3 and inhibitors (AMF, RAD, JNJ, MEH, ECP, MHN, SGI). H stands for hydrogen bond and HP stands for hydrophobic interaction.

| Name of amino acid with residue number |  | AMF | RAD | JNJ | MEH | ECP | MHN | SGI | RAF | BAI | TSH |
| --- | --- | --- | --- | --- | --- | --- | --- | --- | --- | --- | --- |
| Cys | 376 |  |  |  |  |  |  | HP |  | HP | HP |
| Asp | 377 | H |  | HP |  | H |  |  |  | HP | HP |
| Ile | 378 |  | H | HP |  | H |  | HP |  | HP | HP |
| Asp | 395 | H | HP | HP | H | HP | H | HP | HP | HP |  |
| Pro | 396 | HP |  | HP | H | HP | HP | HP | HP |  |  |
| Pro | 397 | HP | HP | HP | HP | HP | HP | HP | HP | HP | HP |
| Trp | 398 |  |  |  | HP |  |  |  | HP |  |  |
| Tyr | 406 | HP | HP | HP | H | HP | HP | HP | HP | HP | HP |
| Gly | 407 | HP | HP |  | HP |  |  |  |  | HP | HP |
| Thr | 408 | HP |  |  | HP |  |  |  |  |  |  |
| Leu | 409 | HP | HP | HP | HP | HP |  | HP |  | HP | HP |
| Thr | 433 |  | HP | HP | HP | HP | HP | HP | H |  |  |
| Glu | 481 |  | HP |  | H |  |  |  | HP |  |  |
| Thr | 510 |  | HP |  | H |  |  | H | H |  |  |
| Ser | 511 | HP | HP | HP | H | HP |  | HP | HP | H |  |
| Lys | 513 |  |  | HP | HP | HP |  |  |  | HP |  |
| Phe | 534 | HP | HP | HP | H | HP | HP | HP | HP | H | HP |
| Arg | 536 | HP |  | HP | HP | HP |  |  |  | HP | HP |
| Gly | 548 |  |  | HP | HP | HP |  | HP |  |  |  |
| Asn | 549 | HP | HP | H | H | HP | HP | HP | HP | HP | HP |
| Gln | 550 | HP |  |  |  |  |  |  |  |  |  |
| Arg | 379 | H |  |  | HP |  |  |  |  |  | HP |
| Gly | 535 |  |  | HP |  |  |  |  |  | HP |  |
| Asn | 539 |  |  |  |  |  |  |  |  | H |  |
| His | 512 |  |  |  |  |  |  |  |  | HP |  |
| His | 538 |  |  |  |  |  |  |  |  | H |  |
| Glu | 532 |  |  |  |  |  |  |  |  | H |  |
| Ile | 400 |  | HP |  |  |  |  | HP | HP |  |  |

It is clear that all inhibitor molecules have occupied the same binding pocket that is considered to be the binding pocket for SAM/SAH/STM2457. But if we zoom into the amino acid residue level one can find significant variation in nature and number of contacts between the amino acid residues and the ligand (Table 2). This piece of information could be useful in designing more potent inhibitors using the docked inhibitors as lead.

### Binding energy calculation

The binding energy of the docked protein-small molecule complex was also calculated by Prodigy (<u>Prodigy Webserver (uu.nl)</u>), a server from Bonvin lab, to compare with the ligand/inhibitor bound crystal structures of METTL3. The binding energy score of the best ten hits obtained from Autodock Vina showed better result than the existing crystal structures (Table 3). It also corroborates the docking score and gives another layer of validation that the best scoring compounds are potentially strong inhibitors of METTL3.

**Table 3:** Binding energy (kcal/mol) Binding energy (Kcal/mole) calculation of the 10 best hits along with substrate, substrate analogue and existing inhibitor (STM2457) bound crystal structure.

| **SAM** | **SAH** | **STM**<br>**2457** | **AMF** | **RAD** | **JNJ** | **MEH** | **ECP** | **MHN** | **SGI** | **RAF** | **BAI** | **TSH** |
| --- | --- | --- | --- | --- | --- | --- | --- | --- | --- | --- | --- | --- |
| -7.2 | -7 | -8.9 | -10.6 | -10.6 | -7.8 | -10.6 | -10.7 | -9.8 | -11 | -10.5 | -9.8 | -8.8 |

### Swiss ADME analysis

From the molecular docking analysis, we have shortlisted 10 compounds with significantly high docking score compared to the non-hydrolysable substrate analogue (SAH) and existing inhibitors like STM2457. The binding energy calculation from Prodigy server depicts similar findings while compared with the substrate / substrate analogue and inhibitor bound crystal structures. All these in-silico analyses are suggesting that AMF, RAD, JNJ, MEH, ECP, MHN, SGI, RAF, BAI, TSH, have the potency to be strong inhibitors against METTL3. But it does not make these compounds potential drug molecules until it passes the Lipinski’s rule of five, which considers molecular weight, solubility, number of hydrogen bond donor, number of hydrogen bond acceptor, partition coefficient between n-octanol and water. Compounds violating Lipinski’s rule of five, fail to score enough to be considered as druglike molecule. These pharmacokinetic properties were estimated using open-source Swiss ADME server (http://www.swissadme.ch/). Determining the pharmacokinetic properties are important in drug discovery because more than 40% of the compounds get rejected due to poor score in ADME [41]. The pharmacokinetic properties of the selected inhibitor molecules are presented in table 4. Out of the ten best hits, seven compounds, RAD, RAF, JNJ, ECP, MHN, TSH, SGI are obeying the Lipinski’s rule are has been found to be druggable. On the other hand, AMF, MEH, BAI are less druglike due to some violations in the Lipinsiki’s rule but it does not undermine the essentiality of these compounds in epitranscriptomics related research. Because other than the drug discovery for cancer therapeutics, finding potent inhibitors against METTL3 is equally important to study the role of METTL3 in different cellular processes as well as in cancer progression. AMF, MEH, BAI also has the potential to be lead compounds that could be chemically modified to enhance the druglikeness.

**Table 4:** ADME prediction. The five major physicochemical parameters such as molecular weight (MW), number of hydrogen bond acceptors (NHBA), number of hydrogen bond donors (NHBD), molar refractivity (MR) and lipophilicity (ilogp) of all the ten hits are shown here. The toxicity score & class has been calculated using Toxi M and Protox II server respectively.

| Parameters | AMF | RAD | JNJ | MEH | ECP | MAH | SGI | RAF | BAI | TSH |
| --- | --- | --- | --- | --- | --- | --- | --- | --- | --- | --- |
| **MW** | 538.46 g/mol | 518.41 g/mol | 365.45 g/mol | 624.59 g/mol | 396.56 g/mol | 347.45 g/mol | 461.52 g/mol | 504.39 g/mol | 446.36 g/mol | 276.29 g/mol |
| **Acceptable range** | $\leq 500$ | $\leq 500$ | $\leq 500$ | $\leq 500$ | $\leq 500$ | $\leq 500$ | $\leq 500$ | $\leq 500$ | $\leq 500$ | $\leq 500$ |
| **NHBA** | 10 | 10 | 3 | 15 | 3 | 2 | 4 | 10 | 11 | 3 |
| **Acceptable range** | $\leq 10$ | $\leq 10$ | $\leq 10$ | $\leq 10$ | $\leq 10$ | $\leq 10$ | $\leq 10$ | $\leq 10$ | $\leq 10$ | $\leq 10$ |
| **NHBD** | 6 | 2 | 1 | 7 | 1 | 2 | 4 | 3 | 6 | 0 |
| **Acceptable range** | $\leq 5$ | $\leq 5$ | $\leq 5$ | $\leq 5$ | $\leq 5$ | $\leq 5$ | $\leq 5$ | $\leq 5$ | $\leq 5$ | $\leq 5$ |
| **MR** | 146.97 | 122.44 | 111.02 | 145.88 | 117.49 | 110.48 | 140 | 117.53 | 106.72 | 80.24 |
| **Acceptable range** | 40–130 | 40–130 | 40–130 | 40–130 | 40–130 | 40–130 | 40–130 | 40–130 | 40–130 | 40–130 |
| **ilogp** | 3.06 | 2.78 | 3.31 | 1.4 | 4.09 | 3.52 | 2.94 | 2.13 | 1.75 | 2.44 |
| **Acceptable range** | $\leq 5$ | $\leq 5$ | $\leq 5$ | $\leq 5$ | $\leq 5$ | $\leq 5$ | $\leq 5$ | $\leq 5$ | $\leq 5$ | $\leq 5$ |
| **Druglikeness** | No (2 Violations) | Yes (1 Violation) | Yes (0 Violation) | No (3 Violations) | Yes (1 Violation) | Yes (0 Violation) | Yes (0 Violation) | Yes (1 Violation) | No (2 Violations) | Yes (0 Violation) |
| **Toxi M score** | 0.92 | 0.929 | 0.989 | 0.791 | 0.716 | 0.959 | 0.921 | 0.912 | 0.706 | 0.98 |
| **ProTox class** | 5 | 4 | 5 | 6 | 5 | 5 | 5 | 4 | 5 | 4 |

The relative toxicity of the selected small molecule inhibitors were also assessed using ToxiM (ToxiM (iiserb.ac.in)) and ProTOX II (ProTox-II - Prediction of TOXicity of chemicals (charite.de))server. It shows that all these ten compounds are belong to toxicity class 4-6 according to ProTOX II server, which is within the acceptable range. This information gives a preliminary understanding about the druggability of these molecules.

### Molecular Dynamics Simulation

The stability of the top 7 protein-ligand complexes derived from molecular docking were studied using molecular dynamics simulation and compared with apo-protein simulation results. It has been found that the overall RMSD of the apo METTL3 and the METTL3-inhibitor complex (Fig 2A) are within the allowable limit (< 0.3 Å) for the globular protein [42]. The average RMSD of the apo-protein is 0.188Å. The overall average RMSD of the proteins bound to different inhibitors is comparable to the average RMSD of the protein alone, ranging from 0.156-0.213Å. The change of RMSD value of the protein backbone for different inhibitor-protein complexes with the apo-protein was compared over course of simulation by plotting the RMSD values against time (Fig 2). The average RMSD of the different ligand bound complexes are tabulated in Table 5. In presence of SGI and RAD the average overall RMSD is lower than the protein alone which signifies that the binding of these two aforementioned ligands stabilizes the METTL3 structure, indicating their interaction with the protein was stable throughout the simulation process.

**Figure 2:**
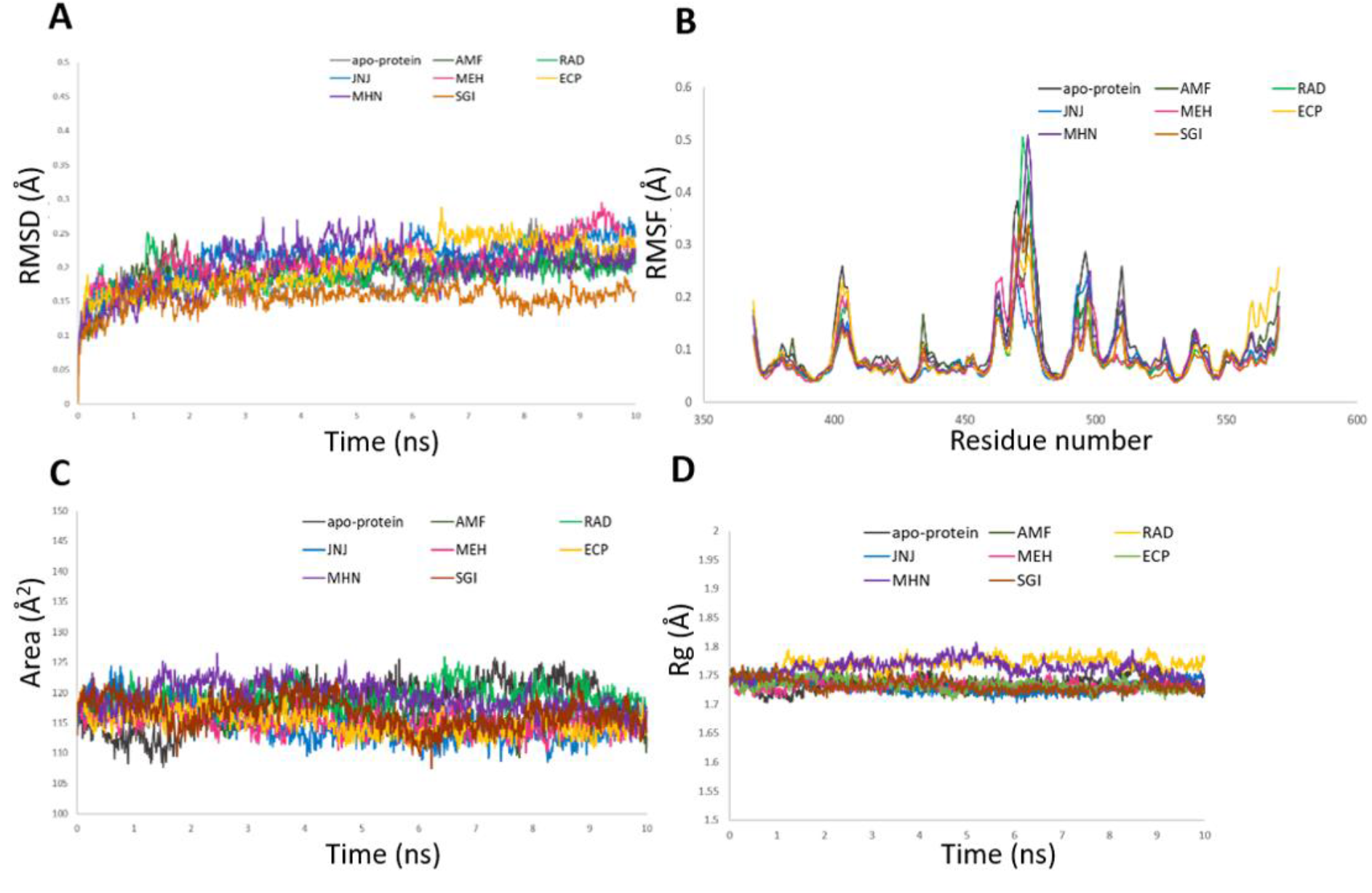
The graphical representation of the MD simulation results of apo-protein and in the presence of inhibitor molecules. AMF, RAD, JNJ, MEH, ECP, MHN, SGI throughout the simulation time frame (10 ns). (A) The RMSD (Å) of Mettl3 backbone in the absence (apo-protein) and presence of inhibitors, (B) Mettl3 backbone RMSF (Å) against the amino residue number, (C) Radius of gyration (Rg) in Å vs time (ns) and (D) solvent accessible surface area (SASA) in Å^2^ vs time (ns).

**Table 5:** Molecular dynamics simulation result. Average Root Mean Square Deviation (RMSD), Root Mean Square Fluctuation (RMSF), Radius of Gyration (Rg), Solvent Accessible Surface Area (SASA) and protein ligand interaction energy (Lennard-Jones energy) has been calculated from MD simulation run.

| **Sr. No.** | **Compound Code** | **Average RMSD (Å)** | **Average RMSF (Å)** | **Average Rg (Å)** | **Average SASA (Å<sup>2</sup>)** | **Protein-ligand interaction energy (kcal/mol)** |
| --- | --- | --- | --- | --- | --- | --- |
| 1 | Apo protein | 0.189 | 0.108 | 1.737 | 118.296 | - |
| 2 | AMF | 0.193 | 0.096 | 1.733 | 116.041 | -42.4 |
| 3 | RAD | 0.188 | 0.089 | 1.769 | 119.128 | -39.6 |
| 4 | JNJ | 0.213 | 0.087 | 1.730 | 114.708 | -39.0 |
| 5 | MEH | 0.207 | 0.087 | 1.732 | 115.104 | -42.3 |
| 6 | ECP | 0.202 | 0.096 | 1.734 | 115.162 | -32.7 |
| 7 | MHN | 0.198 | 0.097 | 1.763 | 119.257 | -34.2 |
| 8 | SGI | 0.156 | 0.083 | 1.732 | 116.563 | -38.2 |

#### RMSF

It calculates the fluctuation of the individual amino acid residues throughout the simulation process in the presence and absence of different ligand molecules (Fig 2B). The RMSF of AMF, SGI, RAD, MHN, MEH, ECP, JNJ are lower than the RMSF of the apo-protein, which signifies ligand binding has stabilized the fluctuation of amino acid sidechains.

#### SASA

The solvent accessible surface area (SASA) of the protein alone is 118 Å^2^. For inhibitor bound complexes, the SASA varies from 115-119 Å^2^. For most inhibitor bound complexes, AMF, SGI, MHN, ECP, JNJ, the SASA is less than that of the protein alone. For RAD and MEH, the SASA is slightly higher (~119 Å^2^), that could be due to a conformational change. RMSF analysis suggests that the amino acids 467-486 have comparatively higher fluctuation for RAD and MEH (Fig 2D).

#### Radius of gyration

The compactness of protein is indicated by radius of gyration (R_g_). The average R_g_ value for the protein is 1.736Å. For MEH, SGI, AMF, JNJ, RAD and ECP the average R_g_ is comparable with the apo-protein. The values are given in Table 5.

## Discussion

The extensive *in-silico* analysis and molecular docking of 810 commercially available small molecules, including natural compounds, are clearly indicating that AMF, RAD, SGI, JNJ, MEH, MHN, ECP could be used as the potent inhibitor of METTL3. Some of these compounds are showing no violation to Lipinski’s rule thus claim to be compelling drug candidates for cancer therapeutics. If compared with the substrate, substrate analogue and inhibitor bound crystal structures of METTL3, the data clearly indicates that these seven compounds are showing significantly better score in docking and in the terms of binding energy calculation. The molecular dynamics simulation output parameters are also in good agreement with the docking results, where it is clearly showing that docked protein-ligand complexes remain stable throughout the entire simulation process. These compounds are potent inhibitors of METTL3 and promising drug molecules against METTL3 induced cancer. Other than being effective drug candidates, these candidates hold much promise in the search of strong inhibitors against METTL3. Discovery of strong inhibitor against METTL3 is immensely important to explore the role of epitranscriptomic modification in different layers of molecular regulation. These small molecules are also providing unique scaffolds to design new more effective compounds in future.

The natural compounds found to be potential inhibitors of METTL3 have therapeutic roles, AMF, a biflavonoid, is known to act as an inhibitor in cancer [43, 44] and malaria [45]. Docking results indicate AMF binds strongly with METTL3 this is validated by simulation studies as well. The AMF-METTL3 complex was more stable during the period of simulation than the protein alone. However, AMF does not qualify the Lipinski’s rule of five. Searching for small molecule derivatives of AMF, we found Bilobetin [46] which also gives a high docking score (−10.5 kcal/mol). RAD is a derivative of RAF265 and found to be VEGFR and RAF kinase inhibitor [47]. JNJ has been found as potent selective inhibitor of fatty acid amide hydrolase (FAAH) and already has been used in developing new drugs against neuropathic pain treatment [48]. The role of METTL3 in AML, CRC, HCC, lung cancer is well established, and inhibiting METTL3 has been proven to be useful to limit cancer progression. Therefore, the compounds found in this study can be highly useful to serve as potential lead molecules to be developed into drugs.

## Methods

### Receptor preparation for docking

The X-ray crystal structure of human METTL3 in the S-Adenosyl Homocysteine (SAH) bound form (PDB ID 5IL2, resolution: 1.61 Å) was retrieved from Protein data bank (https://www.rcsb.org/) as the receptor for molecular docking. The structure consists of methyltransferase domain of METTL3 protein with amino acid boundary 369 to 570. The SAM binding pocket of the receptor was targeted for virtual screening. Prior to docking, the heteroatoms (ligand, water, ions) were removed from the receptor PDB file. Polar hydrogens were added, charges were merged; nonpolar hydrogens and lone pairs were removed and receptor PDBQT file was generated using UCSF Chimera [49]. The SAH (PDB ID 5IL2) and SAM bound (PDB ID 5IL1) crystal structures were used to validate the docking results.

Ligand preparation for virtual screening: 810 commercially available compounds were selected for in silico screening through extensive literature survey and careful analysis of available small molecule libraries like Pubchem (https://pubchem.ncbi.nlm.nih.gov/). The small molecules were converted to the 3D structure, Gasteiger charges were added, and merged; nonpolar hydrogens, lone pairs were removed and ligand PDBQT file generation was done by UCSF Chimera. The list of all the selected compounds is present in the supplementary file.

The binding energy of the small molecules that had best score were rechecked using PRODIGY server from Bonvin Lab.

### Receptor grid generation, molecular docking and binding energy calculation

Docking was performed using Autodock Vina, an opensource software through UCSF Chimera GUI. The search grid was set including the residues of SAM/SAH binding pocket (ASP377, ASP395, ASN539, GLU532, ARG536, HIS538, ASN549, & GLN550). For each run, number of binding mode was 5.0, exhaustiveness of run was 8.0 and maximum energy was 2.0 kcal/mol. Validation is a crucial step in virtual screening. To validate the docking protocol, SAH, the co-crystal ligand of METTL3 from 5il2 was redocked with METTL3 and then the docked complex was superimposed with the co-crystal to compare the binding of SAH. The docking score of SAH was considered as the standard reference. To validate the docking protocol and ensure the small molecules were docked in the SAM binding pocket, the docked structures were superimposed on the native conformation of SAH bound METTL3 receptor. Once the grid box and docking parameters were validated, the selected 810 compounds were subjected for docking using the same protocol. The interaction energy between the binding pocket in the receptor protein and the ligand was calculated from the molecular docking process. The compounds were scored on their ability to interact with the SAM binding pocket of the target protein. The compounds were first screened and the resulting scores were sorted and ranked. A threshold of binding energy −9 kcal/mol was set up to screen only those molecules that have a stronger binding compared to either SAH or the existing commercially available inhibitor of METTL3, STM2457. The top compounds were ranked according to their binding affinities and the ligand–protein binding free energy (ΔG) as well as the list of atoms present within 10.5Å at the interface between protein & ligand were calculated using ProdigY (PROtein binDIng enerGY prediction) server (<u>Prodigy Webserver (uu.nl)</u>).

### Determination of Ki using Autodock 4.2.6

Top 10 compounds, based on the Autodock Vina score were subjected to another round of docking with Autodock 4.2.6 using Autodock tools. The same receptor PDB file was used for docking. Kollman charges were added and the corresponding PDBQT file was prepared using Autodock tools. Search grid box was prepared by targeting the SAM binding pocket as described in the previous section. The ligand PDBQT files were generated via Open Babel software. For the ligands Gasteiger charges have been added prior to docking. For docking search parameter and docking output generation genetic algorithm and Lamarckian genetic algorithm were used respectively in search of the lowest binding energy. The Autodock 4.2.6 score and the corresponding Ki for the docked compound have been tabulated in the table 1.

### Drug like properties and toxicity prediction

To evaluate the druglikeness of the top scoring compounds selected on the basis of docking score, Ki and Prodigy energy were subjected to ADME properties determination using open source SwissADME server. The five physicochemical parameters (Lipinsiki’s rule) that is molecular weight, number of H-bond acceptors (NHBA), number of H-bond donors (NHBD), molar refractivity, n-octanol/water partition coefficient were calculated and tabulated in table 4. The compounds showing promising results were selected for protein-ligand simulation study. The level of toxicity was also calculated using the open source ToxiM and ProTox II server.

### MD simulation protocol

For further evaluation the 7 top performing ligands were subjected to molecular dynamics simulation (MDS) of the receptor protein-ligand complex. The all atom MDS of these selected protein-ligand complexes were performed by Gromacs 5.1.2 [50] Molecular Dynamics packages (https://www.gromacs.org/). The protein topology was prepared by the Charmm36 force field [51]. For each ligand, H atom was added with Avogadro [52] and topology file was generated using The CHARMM General Force Field (CGenFF) server [51]. After compiling the protein and ligand topology the complex topology file was solvated using TIP3P [53] water model in a defined dodecahedron box and rendered electroneutral by the addition of required number of Na^+^ and Cl^-^ counter ions. For energy minimization each system with a maximum number of 50000 steps was done using the steepest descent algorithm and the force was set to less than 10.0 kJ/mol. The two-stage equilibration step consists of the NVT ensemble step and the NPT ensemble step. In the NVT ensemble step the number of particles (N), volume (V) and temperature (T) i.e., 300 K were kept constant and maintained for 100ps. The second step or the NPT ensemble step has constant pressure (P) i.e., one bar along with the equilibration of temperature (T) and number of particles (N) for 100ps. Berendsen’s method [54] and Parrinello-Rahman barostat [55] methods were used to perform NVT and NPT equilibration respectively. Upon the execution of the two equilibrium phases the equilibrated system was subjected to MD run for 10 ns (5000000 steps). On successful completion of the MDS, removal of water and ions was done followed by PBC correction for the purification of the trajectories. These purified trajectories assist to analyse and calculate various parameters such as root mean square deviation (RMSD) [56], root mean square fluctuation (RMSF) [57], radius of gyration (Rg) [58], solvent accessible surface area (SASA) [59] and protein-ligand interaction energy (short range Lennard-Jones energy) [50]. The Chimera 1.15 software was used to visualize the trajectory and render MD movie/clip and images.

## Acknowledgement

This work has been carried out with the support of JIS University, Kolkata. Authors would like to thank Prof. Indranil Sengupta, honourable Vice Chancellor, JIS University for providing the necessary infrastructure for the work. Authors acknowledge Mr. Sumanta Ghosh and Mr. Soumalya Chowdhury from the central computational facility of JIS University for providing technical help. The authors also thank Dr. Mainak Mukhopadhyay for his constant support and encouragement.

## Author contributions

AR and DH conceptualized the work, designed the experiments, performed MD simulation and analysed the data. SM, PS, RB performed docking, pharmacokinetic analysis and compiled the results. AKR, RK and AA performed docking. AM, SM, AR and DH wrote the manuscript.

## Data availability

All raw data files are available.

