## supplemental figure1, 2, supplemental table 1 for "Identification of potential natural compound inhibitors and drug like molecules against human METTL3 by docking and molecular dynamics simulation"

**Supplementary Figure 1:** Superposition of docked SAH (yellow) and SAH (green) bound crystal structure of METTL3 (PDB ID 5IL2).

**Supplementary Figure 2:** LIGPLOT analysis of best 7 small molecule inhibitors. The hydrogen bonds are shown in green dotted line. The hydrophobic interactions are projected as arcs with radiating spokes.

**Supplementary Table 1:** The complete list of 810 commercially available compounds, screened using Autodock Vina as the docking platform along with the ADV score.

**Supplementary Figure 1**

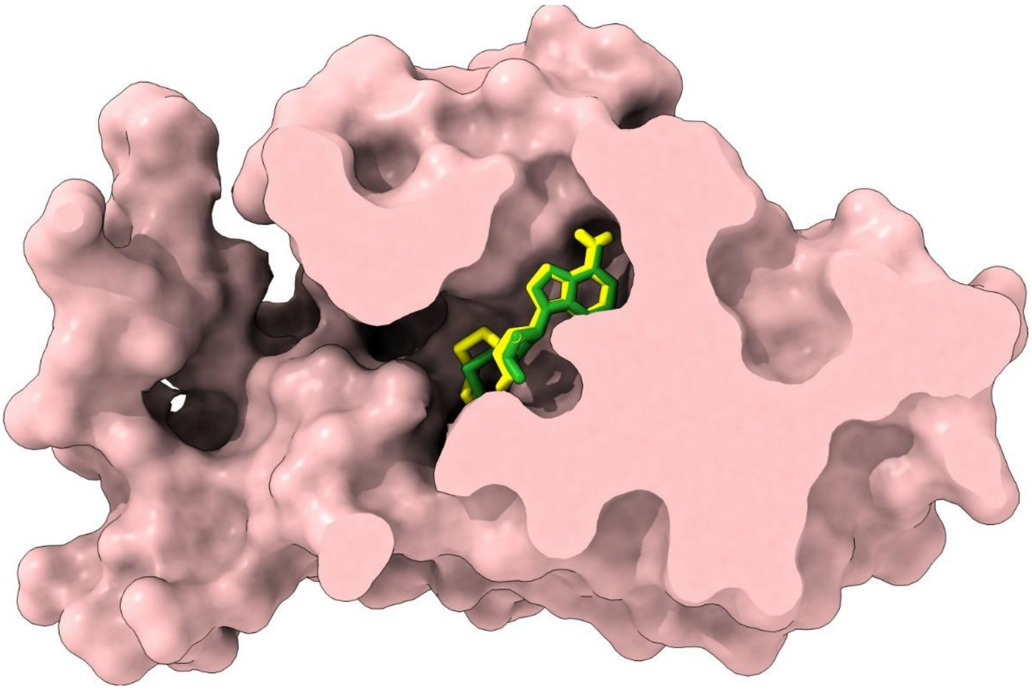

### Supplementary Figure 2

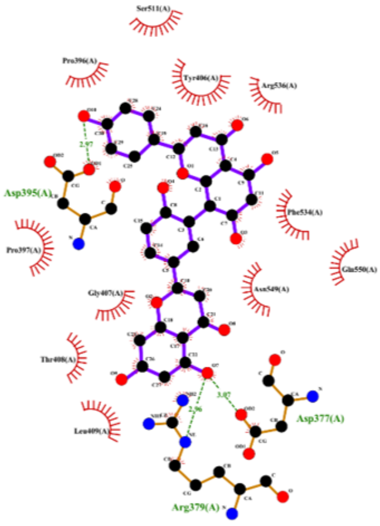

AMF

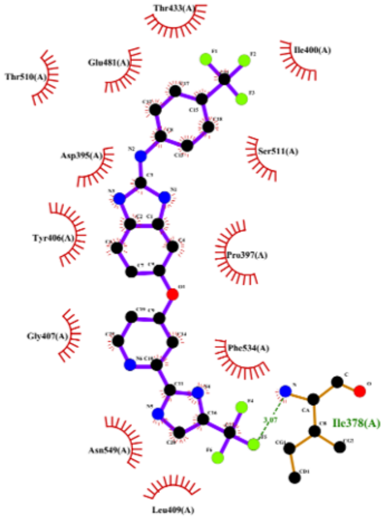

RAD

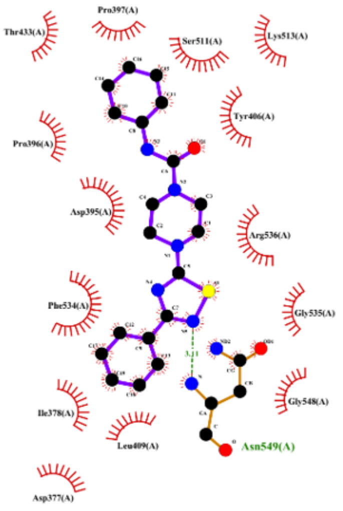

JNJ

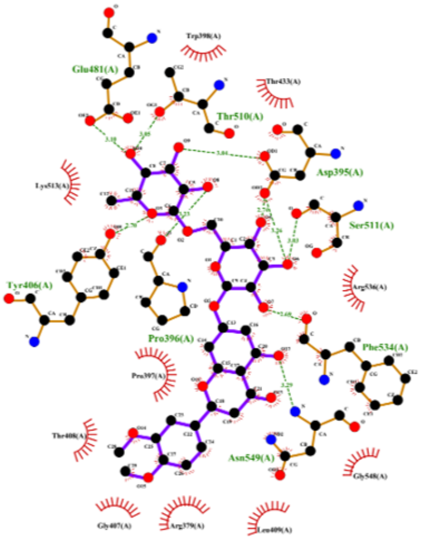

MEH

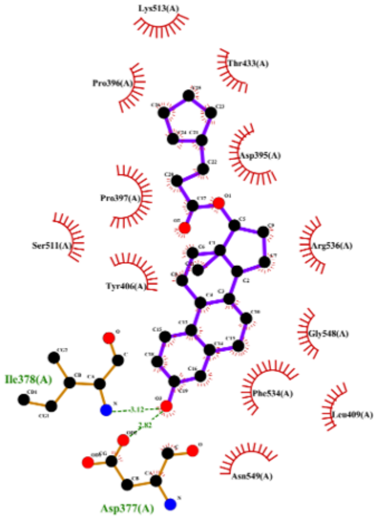

ECP

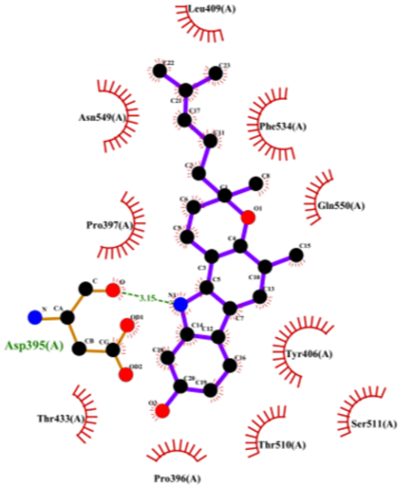

MHN

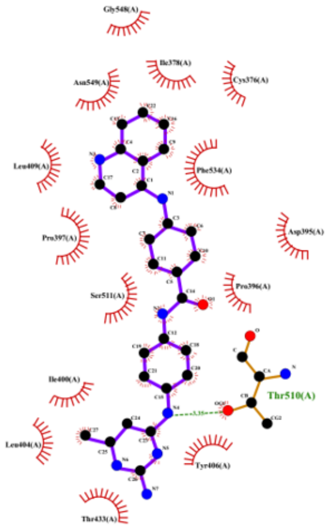

SGI

**Supplementary Table 1**

| Sl No | Name of compound | Vina Score |
|---|---|---|
| 1 | (-)-Borneol | -5.1 |
| 2 | (-)Epicatechin | -8.1 |
| 3 | (-)-Menthol | -5.4 |
| 4 | (-)-Parthenolide | -7.8 |
| 5 | (+)- Usniacin | -7.5 |
| 6 | (+)-Borneol | -5.3 |
| 7 | (+)-Catechin | -8.4 |
| 8 | (+)-Catechin hydrate | -1.7 |
| 9 | (+)-Fangchinoline | -8.5 |
| 10 | (+)-Matrine | -6.8 |
| 11 | (1R,2R)-trans-N-Boc-1,2-cyclohexanediamine | -5.5 |
| 12 | (20S)-Protopanaxadiol | -8.2 |
| 13 | (20S)-Protopanaxatriol | -8.1 |
| 14 | (E)-Cardamoni | -7.8 |
| 15 | (S)-10-Hydroxycamptothecin | -9.5 |
| 16 | 10-deacetylbaccatin-III | -7.4 |
| 17 | 1-Deoxynojirimycin | -5.9 |
| 18 | 2' -DEOXY-5-FLUOROCYTIDINE | -6.4 |
| 19 | 2,4-Dihydroxyacetophenone | -5.6 |
| 20 | 2,6-Dihydroxypurine | -6.4 |
| 21 | 20-hydroxyecdysone | -7.9 |
| 22 | 2'-Deoxyinosine | -7.9 |
| 23 | 2-Methoxy-1,4-naphthoquinone | -7 |
| 24 | 2-Methoxyestradiol (2-MeOE2) | -8.8 |
| 25 | 3,3'-Diindolylmethane | -8.5 |
| 26 | 3,4'-5-TRIMETHOXY TRANS STILBENE | -8.2 |
| 27 | 3,5-Diiodotyrosine Dihydrate | -1.7 |
| 28 | 3-HYDROXY FLAVONE | -8 |
| 29 | 3'-HYDROXYPTEROSITLBENE | -7.7 |
| 30 | 3-Indolebutyric acid (IBA) | -6.4 |
| 31 | 4',7-DIMETHOXY ISO FLAVONE | -8.1 |
| 32 | 4',7-Dimethoxy-5-Hydroxyflavone | -8.7 |
| 33 | 4-Amino-5-imidazolecarboxamide | -5 |
| 34 | 4-BUTYLRESORCINOL | -5.7 |
| 35 | 4-Chloro-2-hydroxybenzoic acid, 4-chloro salicylic acid | -5.7 |
| 36 | 4-Demethylepipodophyllotoxin | -7.6 |
| 37 | 4'-Demethylpodophyllotoxin | -8.3 |
| 38 | 4-Hydroxy-3,5-dimethoxybenzyl alcohol | -5.8 |
| 39 | 4-Hydroxybenzoic acid | -5.5 |
| 40 | 4-Hydroxybenzyl alcohol | -5.3 |

| | | |
|---|---|---|
| 41 | 4-Hydroxychalcone | -7.7 |
| 42 | 4'-METHOXY RESVERATROL | -8 |
| 43 | 4-METHYLESCULETIN | -7.1 |
| 44 | 4-Methylumbelliferon (4-MU) | -7.3 |
| 45 | 5 Fluorozebularine | -6.7 |
| 46 | 5,4'-DIHYDROXY-6,7-DIMETHOXY FLAVONE | -8.1 |
| 47 | 5,7-DIMETHOXY-8-METHTL FLAVONE | -8.2 |
| 48 | 5-Aminolevulinic acid HCl | -4.7 |
| 49 | 5-HYDROXY FLAVONE | -8.3 |
| 50 | 5-Hydroxymethylfurfural | -4.8 |
| 51 | 5-hydroxytryptophan (5-HTP) | -7.1 |
| 52 | 5-methoxyflavone | -8 |
| 53 | 5-Phenyl-2,4-pentadienoic acid | -6.5 |
| 54 | 6-Gingerol | -6.4 |
| 55 | 6-HYDROXY FLAVONE | -8 |
| 56 | 6-Hydroxyflavone (6-HF) | -7.9 |
| 57 | 7,8-DIMETHOXY FLAVONE | -7.7 |
| 58 | 7-HYDROXY FLAVONE | -8.3 |
| 59 | 7-Hydroxy-3,4-dihydrocarbostyril | -6.5 |
| 60 | 7-Methoxy-4-methylcoumarin | -6.9 |
| 61 | 9-Aminoacridine | -7.5 |
| 62 | A 196 | -8.8 |
| 63 | A 366 | -7.7 |
| 64 | ABYSSINONE | -9 |
| 65 | Acarbose | -8 |
| 66 | Acetylcholine iodide | -4.3 |
| 67 | Acetylspiramycin (ASPM) | -6.2 |
| 68 | Acetylvanillin | -5.7 |
| 69 | Adenosine | -7.5 |
| 70 | Adrenosterone | -8.4 |
| 71 | AFZELECHIN | -8.4 |
| 72 | Albendazole | -7.6 |
| 73 | Alizarin | -8.5 |
| 74 | Allantoin | -6.1 |
| 75 | Allicin | -3.2 |
| 76 | Aloe-emodin | -8.4 |
| 77 | Aloin | -8.2 |
| 78 | alpha-Asarone | -5.8 |
| 79 | ALPINETIN | -8.1 |
| 80 | Alvespimycin (17-DMAG) HCl | -6.4 |
| 81 | Amentoflavone | -11.3 |
| 82 | AMI 1 | -10.6 |
| 83 | Aminophyllin | -5.8 |
| 84 | Aminothiazole | -3.7 |
| 85 | Ammonium Glycyrrhizinate | -1.3 |

| | | |
|---|---|---|
| 86 | Amodiquin | -1.7 |
| 87 | Amorolfine HCL | -8 |
| 88 | Amoxicillin | -7.8 |
| 89 | Amoxicillin Sodium | -8.3 |
| 90 | Amphotericin B | -6.6 |
| 91 | Ampicillin Trihydrate | -6.3 |
| 92 | Amygdalin | -6.9 |
| 93 | Andrographolide | -8.4 |
| 94 | ANOMORELIN | -7.2 |
| 95 | ANONAINE | -9.1 |
| 96 | Apigenin | -8.6 |
| 97 | APIGENIN | -8.6 |
| 98 | Apocynin | -5.6 |
| 99 | Arbutin | -7.3 |
| 100 | ARBUTIN | -7.3 |
| 101 | ARCTIGENIN | -8.5 |
| 102 | Arctiin | -8.6 |
| 103 | Arecoline HBr | -4.9 |
| 104 | AROMADENDRIN | -8.3 |
| 105 | ARTEETHER | -7.7 |
| 106 | Artemether | -7.9 |
| 107 | Artemisinin | -8.1 |
| 108 | Artesunate | -8.4 |
| 109 | Asaraldehyde | -5.5 |
| 110 | Asiatic Acid | -6.2 |
| 111 | Asiaticoside | -7.6 |
| 112 | Astaxanthin | -7 |
| 113 | ASTILBIN | -8.4 |
| 114 | astragaloside A | -7.2 |
| 115 | ASTRAGALUS POLYPHENOLS | -8 |
| 116 | Atropin Sulfate Monodrate | -5.8 |
| 117 | ATROPINE | -7.7 |
| 118 | Avibactam sodium | -6.2 |
| 119 | Azacytidine | -7.3 |
| 120 | AZATHRAMYCIN | -5.7 |
| 121 | Azithromycin Dihydrate | -6 |
| 122 | Azomycin | -4.7 |
| 123 | Aztreonam | -7.4 |
| 124 | BACCATIN III | -5.8 |
| 125 | Baicalein | -8.3 |
| 126 | Baicalin | -9.9 |
| 127 | BAICLEIN | -8.3 |
| 128 | Balofloxacin | -6.3 |
| 129 | Baohuoside I | -8.7 |
| 130 | Batyl alcohol | -5.4 |

| | | |
|---|---|---|
| 131 | BAVACHININ | -9.3 |
| 132 | BCI-121 | -7.4 |
| 133 | Benzoic Acid | -6.1 |
| 134 | Benzyl isothiocyanate | -5 |
| 135 | Berbamine (dihydrochloride) | -8 |
| 136 | BERBERINE | -8.3 |
| 137 | Berberine chloride | -1 |
| 138 | Bergapten | -7.2 |
| 139 | Bergenin | -7.9 |
| 140 | Betaine | -3.8 |
| 141 | Betamipron | -6.3 |
| 142 | Betulin | -7.4 |
| 143 | betulinic acid | -7.5 |
| 144 | BETULONIC ACID | -8.5 |
| 145 | Bilobalide | -6.5 |
| 146 | Bilobetin | -10.5 |
| 147 | Bindarit | -8 |
| 148 | BIOCHANIN | -8.2 |
| 149 | Biochanin A | -8.3 |
| 150 | biotin | -6.1 |
| 151 | BIRABRESIB | -8.8 |
| 152 | Bisdemethoxycurcumin (BDMC) | -8.2 |
| 153 | BIX 01294 | -9.9 |
| 154 | BOBCAT 339 HYDROCHLORIDE | -9.4 |
| 155 | BRASSINOLIDE | -8.3 |
| 156 | BRD 4774 | -8.8 |
| 157 | Broxyquinoline | -6 |
| 158 | butenafine hcl | -8.1 |
| 159 | C 7280948 | -7.7 |
| 160 | Cabazitaxel | -6.9 |
| 161 | Caffeic Acid | -6.3 |
| 162 | CAFFEINE | -5.8 |
| 163 | Camphor | -5.2 |
| 164 | Capsaicin(Vanilloid) | -6.3 |
| 165 | Carbadox | -7.4 |
| 166 | CARDAMONIN | -7.7 |
| 167 | Carnosic acid | -8.3 |
| 168 | Carvacrol | -5.9 |
| 169 | Caryophyllene oxide | -6.9 |
| 170 | CATALPOL | -7.4 |
| 171 | CATECHIN | -8.4 |
| 172 | catharanthine | -6.1 |
| 173 | CC 90011 BESYLATE | -8.7 |
| 174 | Cefaclor | -6.9 |
| 175 | Cefaperazone | -9 |

| | | |
|---|---|---|
| 176 | Cefdinir | -7.2 |
| 177 | CEFODIZIME SODIUM | -1.2 |
| 178 | Cefonicid sodium | -1.2 |
| 179 | Cefoxitin sodium | -7.1 |
| 180 | Cefsulodin sodium | -1.2 |
| 181 | Ceftibuten dihydrate | -1.7 |
| 182 | Ceftiofur HCl | -6.2 |
| 183 | Celastrol | -7 |
| 184 | cephalomannine | -6.6 |
| 185 | CEPHALOTOXIN | -8.8 |
| 186 | Cepharanthine | -9 |
| 187 | CHALCONARINGENIN | -7.8 |
| 188 | Chloramphenicol | -7.1 |
| 189 | Chlorhexidine 2HCl | -9.9 |
| 190 | Chlorogenic Acid | -8.3 |
| 191 | Chlorquinaldol | -6.3 |
| 192 | cholecalciferol (Vitamin D3) | -5.9 |
| 193 | Cholic acid | -7.2 |
| 194 | Chrysin | -8.7 |
| 195 | CHRYSIN | -8.7 |
| 196 | chrysophanic acid | -8.3 |
| 197 | Cinchonidine | -8.4 |
| 198 | Cinchonine | -8.5 |
| 199 | Cinnamaldehyde | -5.3 |
| 200 | Cinnamic acid | -5.8 |
| 201 | Cinoxacin | -7.3 |
| 202 | cis-Anethole | -5.4 |
| 203 | Clarithromycin | -7.1 |
| 204 | Clinafoxacin HCl | -8.4 |
| 205 | Clindamycin | -7.5 |
| 206 | Clindamycin HCl | -6.6 |
| 207 | Clofoctol | -7.9 |
| 208 | Clotrimazole | -7.6 |
| 209 | CM 272 | -8.7 |
| 210 | COCAINE | -7.9 |
| 211 | Colchicine | -8.6 |
| 212 | Cordycepin | -7.3 |
| 213 | Corticosterone | -8.2 |
| 214 | Cortisone acetate | -6 |
| 215 | Cortodoxone | -7.4 |
| 216 | Costunolide | -7.3 |
| 217 | Coumarin | -6.5 |
| 218 | CP 2 | -9.1 |
| 219 | CPI 0610 | -8.9 |
| 220 | CPI 169 | -9.2 |

| | | |
|---|---|---|
| 221 | CPI 360 | -9.6 |
| 222 | CPI455 HCL | -7.8 |
| 223 | Crocin | -8.5 |
| 224 | Cryptotanshinone | -9.7 |
| 225 | CURCUMENOL | -7.2 |
| 226 | Curcumin | -8.5 |
| 227 | curcumol | -7.1 |
| 228 | CYANIDIN | -8.6 |
| 229 | CYCLEN | -4.8 |
| 230 | CYCLOASTRAGENOL | -7.8 |
| 231 | Cylcocytidine Hcl | -6.5 |
| 232 | Cysteamine HCl | -2.5 |
| 233 | Cystine | -6.1 |
| 234 | Cytidine | -6.2 |
| 235 | CYTOSINE | -4.9 |
| 236 | D panthenol | -5.6 |
| 237 | D-(+)-Trehalose dihydrate | -1.7 |
| 238 | DAIDZEIN | -8 |
| 239 | DAIDZIN | -8.9 |
| 240 | Daidzin | -8.9 |
| 241 | DAMINOZIDE | -5 |
| 242 | Daphnetin | -7.2 |
| 243 | Daunorubicin HCl | -8.6 |
| 244 | Decitabine | -6.7 |
| 245 | Decursinol angelate | -9 |
| 246 | Dehydroandrographolide Succinate Potasium Salt | -9.7 |
| 247 | Dehydrocostus Lactone | -8.3 |
| 248 | Dehydroepiandrosterone (DHEA) | -7.8 |
| 249 | Dehydroevodiamine hydrochloride | -9.1 |
| 250 | DELPHINIDINE | -1 |
| 251 | Demeclocycline HCl | -9.3 |
| 252 | Demethylzeylasteral (T-96) | -6.6 |
| 253 | Deoxyarbutin | -6 |
| 254 | Deoxycholic acid | -7.4 |
| 255 | Deoxycorticosterone acetate | -8.2 |
| 256 | Dexamethasone (DHAP) | -8.6 |
| 257 | Dexamethasone Acetate | -8.4 |
| 258 | Dextrose | -5.9 |
| 259 | D-Galactose | -6.1 |
| 260 | Diacerein | -8.6 |
| 261 | Diammonium Glycyrrhizinate | -1.3 |
| 262 | Dicoumarol | -8.8 |
| 263 | DIFLOXACIN | -8.3 |
| 264 | Dihydro artemisinin | -7.8 |
| 265 | Dihydroactinidiolide | -5.9 |

| | | |
|---|---|---|
| 266 | dihydromyricetin | -8.3 |
| 267 | Dihydrotestosterone(DHT) | -7.7 |
| 268 | Dihydrothymine | -4.9 |
| 269 | Dinitolmide | -6.2 |
| 270 | DIOSMETIN | -8.7 |
| 271 | Diosmetin | -8.7 |
| 272 | Dirithromycin | -6 |
| 273 | DL- Carnitine-HCL | -4.6 |
| 274 | D-Mannitol | -5.5 |
| 275 | Docetaxel | -7.1 |
| 276 | Dopamine HCl | -6 |
| 277 | Doripenem Hydrate | -1.7 |
| 278 | Doxorubicin (Adriamycin) HCl | -7.2 |
| 279 | Doxycycline Hyclate | -2.6 |
| 280 | D-Pinitol | -4.8 |
| 281 | Dulcitol | -4.9 |
| 282 | EBI 2511 | -9.8 |
| 283 | Ecabet sodium | -1.2 |
| 284 | ECHIMIDINE | -6.9 |
| 285 | Echinacoside | -9.6 |
| 286 | Echinocystic acid | -7.7 |
| 287 | Econazole nitrate | -7.4 |
| 288 | EI 1 | -8.4 |
| 289 | Eleutheroside B | -7.5 |
| 290 | Emodin | -8.9 |
| 291 | Enoxolone | -6.4 |
| 292 | ENTACAPONE | -7.4 |
| 293 | EPIAFZELECHIN | -8.2 |
| 294 | Epiandrosterone | -7.6 |
| 295 | EPICATECHIN | -8.1 |
| 296 | EPIFISETINIDOL | -9.1 |
| 297 | Epigallocatechin gallate(EGCG) | -9.1 |
| 298 | Epinephrine bitartrate | -5 |
| 299 | Epirubicin HCl | -8.8 |
| 300 | Epothilone A | -8.1 |
| 301 | EPZ 004777 | -9.4 |
| 302 | EPZ 005687 | -10.5 |
| 303 | EPZ 011989 | -8.9 |
| 304 | EPZ 015666 | -9.1 |
| 305 | EPZ 020411 | -7.5 |
| 306 | Equol | -7.9 |
| 307 | ERIODICTYOL | -8.3 |
| 308 | Erythritol | -4.5 |
| 309 | Erythromycin | -5.8 |
| 310 | Erythromycin Ethylsuccinate | -6.5 |

| | | |
|---|---|---|
| 311 | Esculetin | -6.7 |
| 312 | Esculin | -7.4 |
| 313 | Estradiol Benzoate | -8.9 |
| 314 | Estradiol Cypionate | -10.2 |
| 315 | Estriol | -8.5 |
| 316 | Estrone | -8.9 |
| 317 | Ethambutol 2HCl | -4.5 |
| 318 | Ethyl ferulate | -5.8 |
| 319 | Ethyl Vanillate | -5.7 |
| 320 | Etoposide | -9.4 |
| 321 | Eugenol | -5.4 |
| 322 | Eupatilin | -8.4 |
| 323 | Euphorbiasteroid | -5.7 |
| 324 | Fangchinoline | -8.5 |
| 325 | FARNESOL | -6.1 |
| 326 | Faropenem Sodium | -1.2 |
| 327 | FB 23 | -8.1 |
| 328 | FB 23-2 | -8.2 |
| 329 | Ferulicacid | -6.3 |
| 330 | Fidaxomicin | -7.3 |
| 331 | FISETIN | -8.7 |
| 332 | Fisetin | -8.7 |
| 333 | FISETINIDOL | -8.3 |
| 334 | Flavanone | -8.1 |
| 335 | Flavone | -8.1 |
| 336 | FLAVONE | -8.1 |
| 337 | Florfenicol | -7.2 |
| 338 | Flucloxacillin sodium | -1.2 |
| 339 | Fluconazole | -7.1 |
| 340 | Flucytosine | -5.2 |
| 341 | Fluorouracil (5-Fluoracil, 5-FU) | -5.5 |
| 342 | FORMONONETIN | -8 |
| 343 | Formononetin | -8 |
| 344 | Forskolin | -7 |
| 345 | FORSYTHIN | -8.4 |
| 346 | Fosfomycin Tromethamine | -4.5 |
| 347 | Furaltadone HCl | -8.1 |
| 348 | Fusidine | -7.6 |
| 349 | GALANGIN | -8.7 |
| 350 | GALANTHAMINE | -7.2 |
| 351 | Galanthamine | -6 |
| 352 | Gallic acid trimethyl ether | -5.4 |
| 353 | GALLOCATECHIN | -8.1 |
| 354 | GAMBOGENIC ACID | -8.8 |
| 355 | Gambogic acid | -9.4 |

| | | |
|---|---|---|
| 356 | GAMMA-ORYZANOL | -8.9 |
| 357 | Ganoderic acid A 471002 | -8.1 |
| 358 | Gastrodin | -7.3 |
| 359 | Gatifloxacin | -9.3 |
| 360 | GEISSOPERMINE | -7.1 |
| 361 | Geldanamycin | -6.2 |
| 362 | genipin | -6.4 |
| 363 | geniposide | -5.6 |
| 364 | geniposidic acid | -7.3 |
| 365 | Genistein | -8.3 |
| 366 | Genistein | -7.9 |
| 367 | GENISTIN | -8.1 |
| 368 | Genistine | -8.8 |
| 369 | GENKWANIN | -8.5 |
| 370 | Gentiopicroside | -6.2 |
| 371 | Gentisic acid | -5.8 |
| 372 | Gibberellic acid | -7.8 |
| 373 | Ginkgolide A | -7.6 |
| 374 | Ginkgolide C | -6.7 |
| 375 | Glabridin | -9.3 |
| 376 | Glucosamine hydrochloride | -6 |
| 377 | GLYCITEIN | -8.1 |
| 378 | Glycitin | -8.7 |
| 379 | GLYCOHOLIC ACID | -8.2 |
| 380 | Gossypol acetic acid | -9.3 |
| 381 | gramine | -5.7 |
| 382 | grape seed extract | -7 |
| 383 | Griseofulvin | -7.1 |
| 384 | GSK 126 | -10.3 |
| 385 | GSK 3326595 | -8.9 |
| 386 | GSK 343 | -10.3 |
| 387 | GSK 503 | -10.9 |
| 388 | GSK 591 | -9.2 |
| 389 | GSK 8879552 | -8.8 |
| 390 | GSK J1 | -9.8 |
| 391 | GSK J4 | -9.4 |
| 392 | GSKLSD 1 DIHYDRO CHLORIDE | -6.9 |
| 393 | Guaiazulene | -7.3 |
| 394 | Guanosine | -7.3 |
| 395 | Guggulsterone E&Z | -7.8 |
| 396 | gynostemma extract | -8 |
| 397 | Harmaline | -7.3 |
| 398 | Harmine | -7 |
| 399 | HCL 61 | -8.9 |
| 400 | HEDERAGENIN | -7.9 |

| | | |
|---|---|---|
| 401 | Helicide | -7.2 |
| 402 | Hematoxylin | -8 |
| 403 | HEMIHYDRO CHLORIDE | -1.3 |
| 404 | hesperetin | -8.1 |
| 405 | hesperidin | -8.3 |
| 406 | HESPERITIN | -8.3 |
| 407 | Higenamine hydrochloride | -8 |
| 408 | Histamine | -4.4 |
| 409 | Homoveratrumic acid | -5.9 |
| 410 | honokiol | -7 |
| 411 | Hordenine | -5.5 |
| 412 | HUPERZINE - A | -8.1 |
| 413 | Hydralazine | -6.5 |
| 414 | Hydrocortisone | -7.8 |
| 415 | HYDROXYCOMPTOTHECIN | -7.1 |
| 416 | Hydroxytyrosol | -5.9 |
| 417 | Hydroxytyrosol Acetate | -6.1 |
| 418 | hyodeoxycholic acid (HDCA) | -9.3 |
| 419 | Hyoscyamine | -6.1 |
| 420 | Icariin | -8.5 |
| 421 | Idebenone | -6.1 |
| 422 | INDIGO | -8.8 |
| 423 | Indirubin | -9.2 |
| 424 | Indole-3-carbinol | -5.7 |
| 425 | INDOLE-3-CARBOXYLIC ACID | -6.2 |
| 426 | Inosine | -7.4 |
| 427 | IOX 1 | -6.6 |
| 428 | ipriflavone (osteofix) | -7.4 |
| 429 | Isatin | -5.9 |
| 430 | Isoalantolactone | -7.8 |
| 431 | Isoferulic Acid | -5.4 |
| 432 | ISOHAMNETIN | -9.4 |
| 433 | ISOIMPERATORIN | -7.6 |
| 434 | isoliquiritigenin | -8.1 |
| 435 | ISOPSORALEN | -7.1 |
| 436 | Isoquercitrin | -8.5 |
| 437 | ISOSTEVIOL | -7 |
| 438 | Isotretinoin | -7.9 |
| 439 | Isovaleramide | -4.7 |
| 440 | JATRORRHIZINE | -8.6 |
| 441 | Jervine | -8.6 |
| 442 | JIB 04 | -7.7 |
| 443 | JNJ-1661010 | -10.5 |
| 444 | JQ1 | -7.8 |
| 445 | JQEZ 5 | -9.7 |

| | | |
|---|---|---|
| 446 | KAEMPFERIDE | -8.7 |
| 447 | KAEMPFEROL | -8.6 |
| 448 | kaempferol | -8.5 |
| 449 | Ketoisophorone | -5.4 |
| 450 | kinetin | -5.9 |
| 451 | Kitasamycin | -5.4 |
| 452 | Kynurenic acid | -7.1 |
| 453 | L(+)-Arabinose | -5.5 |
| 454 | L-(+)-rhamnose monohydrate | -4.5 |
| 455 | L-5-Hydroxytryptophan | -7.1 |
| 456 | Lactulose | -6.7 |
| 457 | LADENEIN | -8.6 |
| 458 | lappaconitine | -7.1 |
| 459 | LATHYROL | -7.6 |
| 460 | Lauric Acid | -4.8 |
| 461 | Laurocapram | -5.3 |
| 462 | Lawsone | -7.2 |
| 463 | L-carnitine | -4.5 |
| 464 | Levofloxacin | -8.7 |
| 465 | Ligustrazine hydrochloride | -5.3 |
| 466 | limonin | -8.1 |
| 467 | LINALOOL | -4.9 |
| 468 | Lincomycin HCl | -7.4 |
| 469 | Linezolid | -8.5 |
| 470 | LIQUIRITGENIN | -8.1 |
| 471 | LIQUIRITIN | -9 |
| 472 | LIRAMETOSTAT | -9.7 |
| 473 | Lithocholic acid | -6.7 |
| 474 | LLY 283 | -8.7 |
| 475 | LLY 507 | -7.7 |
| 476 | Lobeline hydrochloride | -8.6 |
| 477 | Loganin | -8.2 |
| 478 | Lovastatin | -5.9 |
| 479 | L-Thyroxine | -7.4 |
| 480 | L-Tryptophan | -6.6 |
| 481 | LUPANINE | -7.5 |
| 482 | LUTEOLIN | -8.8 |
| 483 | luteolin | -8.7 |
| 484 | LYCORINE | -7.8 |
| 485 | Lycorine hydrochloride | -7.6 |
| 486 | LYSIONOTIN | -7.7 |
| 487 | magnolol | -7 |
| 488 | Mahanine | -10.1 |
| 489 | MALTITOL | -6.2 |
| 490 | MALVIDIN | -8.3 |

| | | |
|---|---|---|
| 491 | Meclocycline Sulfosalicylate | -8.4 |
| 492 | MECLOFENAMATE SODIUM | -7.7 |
| 493 | Melatonin | -6 |
| 494 | Melezitose | -6.2 |
| 495 | Melibiose | -7.5 |
| 496 | Menadiol Diacetate | -7.5 |
| 497 | Mequinol | -4.6 |
| 498 | Meropenem | -7.5 |
| 499 | Meropenem Trihydrate | -1.7 |
| 500 | MESTEROLONE | -7.6 |
| 501 | Methoxsalen | -7.4 |
| 502 | Methyl 4-hydroxybenzoate | -5.7 |
| 503 | Methyl 4-hydroxycinnamate | -5.9 |
| 504 | Methyl Cholate | -7.2 |
| 505 | Methyl EudesMate | -5 |
| 506 | Methyl gallate | -6.2 |
| 507 | Methyl protocatechuate | -5.6 |
| 508 | Methyl salicylate | -5.4 |
| 509 | Methyl syringate | -5.7 |
| 510 | Methyl Vanillate | -5.5 |
| 511 | Methyl-Hesperidin | -10.3 |
| 512 | Methylmalonate | -4.6 |
| 513 | Methylnonylketone | -4.4 |
| 514 | Mevastatin | -8.2 |
| 515 | MI 136 | -9.2 |
| 516 | MI 2 | -8.2 |
| 517 | MI 3 | -8.4 |
| 518 | MI 463 | -9.6 |
| 519 | MI 503 | -9.2 |
| 520 | Miconazole | -7 |
| 521 | Miconazole Nitrate | -3.4 |
| 522 | ML 324 | -8.9 |
| 523 | MM 102 | -8.1 |
| 524 | MORIN | -8.7 |
| 525 | Morin Hydrate | -1.7 |
| 526 | MORPHINE | -7.5 |
| 527 | Moxidectin | -7.7 |
| 528 | MS 023 | -6.6 |
| 529 | Mupirocin | -6.5 |
| 530 | Mycophenolic acid | -7.8 |
| 531 | Myricetin | -8.3 |
| 532 | MYRICETIN | -8.3 |
| 533 | Myricitrin | -8.8 |
| 534 | N6-methyladenosine | -6.2 |
| 535 | Nadifloxacin | -8.3 |

| | | |
|---|---|---|
| 536 | Nalidixic acid | -7 |
| 537 | Nanomycin A | -8.4 |
| 538 | NARINGENIN | -8.3 |
| 539 | Naringenin | -8.3 |
| 540 | Naringin | -9.8 |
| 541 | Naringin Dihydrochalcone | -9.3 |
| 542 | Natamycin | -7.3 |
| 543 | Neohesperidin | -9.7 |
| 544 | Neohesperidin Dihydrochalcone (Nhdc) | -8.8 |
| 545 | Neomangiferin | -9.1 |
| 546 | N-Ethylmaleimide (NEM) | -4.4 |
| 547 | Nicotinamide N-oxide | -5.4 |
| 548 | NICOTINE | -5.5 |
| 549 | Nifuratel | -7 |
| 550 | Nithiamide | -5.5 |
| 551 | Nitrofural | -6.2 |
| 552 | Nobiletin | -7.8 |
| 553 | Nocodazole | -8.8 |
| 554 | Noradrenaline bitartrate monohydrate | -1.7 |
| 555 | Norcantharidin | -5.8 |
| 556 | Nordihydroguaiaretic acid (NDGA) | -8.3 |
| 557 | Notoginsenoside R1 | -7.9 |
| 558 | Novobiocin sodium | -1.2 |
| 559 | N-Sulfo-glucosamine sodium salt | -1.2 |
| 560 | Nystatin | -6.6 |
| 561 | Obacunone | -7.9 |
| 562 | octopamine hcl | -5.7 |
| 563 | OG- L002 HCL | -8.3 |
| 564 | OICR - 9429 | -10 |
| 565 | Oleanolic Acid | -7.8 |
| 566 | Oleic Acid | -5.1 |
| 567 | ONAMETOSTAT | -10.9 |
| 568 | Orcinol glucoside | -7.9 |
| 569 | Oridonin | -7 |
| 570 | Ornidazole | -5.3 |
| 571 | Orotic acid (6-Carboxyuracil) | -6.5 |
| 572 | ORY 1001 | -7.5 |
| 573 | Osthole | -7.1 |
| 574 | o-Veratraldehyde | -5.1 |
| 575 | Oxaceprol | -5.9 |
| 576 | Oxymatrine | -7.2 |
| 577 | Oxyresveratrol | -7.8 |
| 578 | Oxytetracycline Dihydrate | -7 |
| 579 | PA-824 | -8.1 |
| 580 | Paclitaxel | -7.7 |

| | | |
|---|---|---|
| 581 | Paeoniflorin | -7.8 |
| 582 | Paeonol | -5.6 |
| 583 | Palmatine | -7.8 |
| 584 | Palmitic acid | -5.3 |
| 585 | Palmitoylethanolamide | -5.2 |
| 586 | Panaxatriol | -6.6 |
| 587 | PAPAVERINE | -8 |
| 588 | Parthenolide | -8 |
| 589 | Patchouli alcohol | -7.7 |
| 590 | Pazufloxacin mesylate | -7.9 |
| 591 | p-Coumaric Acid | -6 |
| 592 | PELARGONIDIN | -8.6 |
| 593 | Pentamidine isethionate | -7.8 |
| 594 | PEONIDIN | -8.7 |
| 595 | Perillyl alcohol | -5.9 |
| 596 | PETUNIDIN | -1 |
| 597 | PF 06726304 | -9.1 |
| 598 | PFI 2HCL | -10 |
| 599 | Phenethyl alcohol | -5.1 |
| 600 | Phloracetophenone | -5.8 |
| 601 | Phloretic acid | -6.2 |
| 602 | Phloretin | -7.8 |
| 603 | PHLORETIN | -7.8 |
| 604 | PHLORIDZIN | -8 |
| 605 | Phlorizin | -8 |
| 606 | Phthalylsulfacetamide | -8.4 |
| 607 | Piceatannol | -8 |
| 608 | Picroside I | -8.7 |
| 609 | Picroside II | -6.7 |
| 610 | Pilocarpine HCl | -5.6 |
| 611 | PINOCEMBRIN | -8.3 |
| 612 | Pinometostat | -10.4 |
| 613 | Piperine | -8.2 |
| 614 | PIPERINE | -8.1 |
| 615 | Piromidic Acid | -8.1 |
| 616 | Plumbagin | -6.9 |
| 617 | Polydatin | -9.6 |
| 618 | Pregnenolone | -8.1 |
| 619 | Primaquine Diphosphate | -6.4 |
| 620 | PROCAINAMIDE HCL | -6.3 |
| 621 | Progesterone | -8.2 |
| 622 | Prostaglandin E2 (PGE2) | -6.5 |
| 623 | Protocatechuic acid | -6.1 |
| 624 | PROTOPINE | -7.8 |
| 625 | Psoralen | -7.4 |

| | | |
|---|---|---|
| 626 | PTEROSTILBENE | -7.5 |
| 627 | Puerarin | -9.7 |
| 628 | PYRIDOXAL | -1.7 |
| 629 | Pyridoxine | -5.3 |
| 630 | Quercetin | -8.6 |
| 631 | Quercetin | -8.7 |
| 632 | QUERCETIN 3,7,3',4'- TETRAMETHYL ETHER | -5.4 |
| 633 | Quercetin Dihydrate | -1.7 |
| 634 | Quercitrin | -8.9 |
| 635 | QUININE | -7.7 |
| 636 | Quinine HCl Dihydrate | -1.7 |
| 637 | Quinolinic acid | -5.5 |
| 638 | Rapamycin (Sirolimus) | -8.9 |
| 639 | Rebaudioside A | -7.6 |
| 640 | Reserpine | -7.7 |
| 641 | Resveratrol | -7.8 |
| 642 | Retapamulin | -6.5 |
| 643 | RG108 | -9.5 |
| 644 | RHEIC ACID | -8.8 |
| 645 | Ribitol | -4.9 |
| 646 | ROBINETINIDOL | -8.4 |
| 647 | Rosavin | -9.3 |
| 648 | Rosmarinic acid | -8.7 |
| 649 | Rotenone (Barbasco) | -8.8 |
| 650 | Rotundine | -8.2 |
| 651 | Roxithromycin | -4.8 |
| 652 | Royal jelly acid | -5.6 |
| 653 | Rutaecarpine | -9 |
| 654 | Salfanilamide | -5.6 |
| 655 | Salicin | -7.2 |
| 656 | Salidroside | -7.7 |
| 657 | SALIRASIB | -8.8 |
| 658 | SALSOLINE | -6.3 |
| 659 | Schisandrin A | -6.7 |
| 660 | schisandrin B | -6.9 |
| 661 | SCHISANTHERIN A | -7.1 |
| 662 | Schizandrol A | -6.9 |
| 663 | Sclareol | -6.5 |
| 664 | Sclareolide | -7.6 |
| 665 | Scoparone | -6.6 |
| 666 | Scopine | -5.5 |
| 667 | Scopolamine HBr | -6.4 |
| 668 | SCOPOLETIN | -6.9 |
| 669 | SECLIDEMSTAT | -8.5 |
| 670 | Secoisolariciresinol diglucoside | -8.1 |

| | | |
|---|---|---|
| 671 | Sennoside B | -7.7 |
| 672 | Serotonin HCl | -6.2 |
| 673 | Sesamin | -9.3 |
| 674 | Sesamol | -5.5 |
| 675 | SGC 0946 | -9.5 |
| 676 | SGC 2085 | -8.2 |
| 677 | SGC 707 | -8.5 |
| 678 | SGI1027 | -11.2 |
| 679 | SGI1027 ANALOGUE | -8.5 |
| 680 | SGI110 | -10.3 |
| 681 | Shikimic acid | -5.9 |
| 682 | Silibinin | -8.8 |
| 683 | Silymarin | -8.6 |
| 684 | Sinapinic Acid | -6.2 |
| 685 | Sinefungin A9145 | -8.9 |
| 686 | Sinomenine | -6.7 |
| 687 | SNDX 5613 | -8.6 |
| 688 | SOLANINE | -7.3 |
| 689 | sophocarpine | -7.1 |
| 690 | Sophoricoside | -8.7 |
| 691 | Sorbitol | -5.7 |
| 692 | SP 2509 | -9.7 |
| 693 | Spectinomycin 2HCl | -6 |
| 694 | Spermidine trihydrochloride | -4.2 |
| 695 | Spiramycin | -6.1 |
| 696 | Stachydrine | -5.1 |
| 697 | Streptozotocin (STZ) | -6.5 |
| 698 | Succinic acid | -4.8 |
| 699 | Sucralose | -6.2 |
| 700 | Sulbactam | -5.7 |
| 701 | Sulfachloropyridazine | -7.5 |
| 702 | Sulfaguanidine | -6.3 |
| 703 | sulfamerazine | -5.7 |
| 704 | Sulfameter | -7.1 |
| 705 | sulfamethazine | -7 |
| 706 | Sulfamethizole | -6.9 |
| 707 | Sulfamethoxazole | -7 |
| 708 | Sulfamethoxypyridazine | -6.8 |
| 709 | Sulfamonomethoxine | -5.5 |
| 710 | Sulfathiazole | -6.6 |
| 711 | Sulfisoxazole | -7.7 |
| 712 | Sulphadimethoxine | -7.4 |
| 713 | SWERTIAMARIN | -7.9 |
| 714 | Synephrine | -5.6 |
| 715 | Synephrine HCl | -5.7 |

| | | |
|---|---|---|
| 716 | Syringaldehyde | -5.7 |
| 717 | Syringic acid | -5.8 |
| 718 | T 3775440 HCL | -7.8 |
| 719 | Tacrine | -1.7 |
| 720 | Tangeretin | -7.4 |
| 721 | TANGERETIN | -7.4 |
| 722 | Tanshinone I | -9.8 |
| 723 | Tanshinone IIA | -9.7 |
| 724 | Taurochenodeoxycholic acid | -7.3 |
| 725 | Tauroursodeoxycholic Acid (TUDCA) | -8.4 |
| 726 | TAXIFOLIN | -8.5 |
| 727 | Taxifolin (Dihydroquercetin) | -8.5 |
| 728 | TAZEMETOSTAT | -9.9 |
| 729 | Tazobactam | -6.4 |
| 730 | TCE 5003 | -8.6 |
| 731 | Tebipenem Pivoxil | -6.7 |
| 732 | Tectoridin | -8.6 |
| 733 | Testosterone Enanthate | -8.2 |
| 734 | TETRA HYDROCURCUMIN | -7.7 |
| 735 | Tetracycline HCl | -8.2 |
| 736 | Tetrahydropalmatine hydrochloride | -7.7 |
| 737 | Tetrahydropapaverine HCl | -8.4 |
| 738 | THEAFLAVIN | -9.6 |
| 739 | Theophylline-7-acetic acid | -6.5 |
| 740 | Thiamphenicol | -7.4 |
| 741 | Thioctic acid | -4.8 |
| 742 | THIOGUANINE | -5.8 |
| 743 | Thioisonicotinamide | -8.7 |
| 744 | Thymopentin | -7.8 |
| 745 | Thymoquinone | -6.3 |
| 746 | TIAMULIN | -5.8 |
| 747 | TIAMULIN FUMARATE | -5 |
| 748 | Tigecycline | -8.3 |
| 749 | Tiglic acid | -4.5 |
| 750 | Tilmicosin | -7.1 |
| 751 | Tinidazole | -4.7 |
| 752 | Tioconazole | -7.1 |
| 753 | TOLCAPONE | -7.9 |
| 754 | Trans-Anethole | -5.5 |
| 755 | Tretinoin | -7.8 |
| 756 | TRIACETONAMINE | -5.4 |
| 757 | TRICIN | -8.5 |
| 758 | trimethoprim | -6.3 |
| 759 | Trimethylaurintricarboxylic acid | -8 |
| 760 | triptolide | -6.7 |

| | | |
|---|---|---|
| 761 | TROPINE | -4.8 |
| 762 | Troxerutin | -9.1 |
| 763 | Tryptamine | -5.7 |
| 764 | Tryptophol | -7.1 |
| 765 | tylosin tartrate | -6.2 |
| 766 | Tyramine | -5.5 |
| 767 | Tyrosol | -5.6 |
| 768 | Umbelliferone | -6.7 |
| 769 | UNC 0379 | -7.7 |
| 770 | UNC 1999 | -10 |
| 771 | UNCO 642 | -7.9 |
| 772 | UNCO 646 | -9.8 |
| 773 | Uracil | -4.8 |
| 774 | Urea | -3.5 |
| 775 | Uridine | -6.6 |
| 776 | UROSONIC ACID | -8.1 |
| 777 | Ursodial | -8.6 |
| 778 | Ursolic Acid | -9.6 |
| 779 | VALEMETOSTAT | -8.7 |
| 780 | Valnemulin HCl | -6.5 |
| 781 | Vanillin | -5.5 |
| 782 | Vanillyl Alcohol | -5.5 |
| 783 | Vanillyl Butyl Ether | -5.4 |
| 784 | Vanillylacetone | -5.9 |
| 785 | VERATRAMINE | -9.3 |
| 786 | Veratric acid | -6 |
| 787 | VINBLASTINE | -5.8 |
| 788 | VINCRISTINE | -5.1 |
| 789 | Vindoline | -5.1 |
| 790 | Vinorelbine Tartrate | -7.3 |
| 791 | Vitamin A Acetate | -7.6 |
| 792 | Vitamin C | -6.1 |
| 793 | Vitamin D2 | -6.5 |
| 794 | Vitamin E | -8.3 |
| 795 | Vitamin E Acetate | -7 |
| 796 | Vitamin K1 | -8.2 |
| 797 | Voriconazole | -7.6 |
| 798 | VTP 50469 | -8.7 |
| 799 | WDR 50103 | -8.2 |
| 800 | Wogonin | -8.6 |
| 801 | Xanthone | -7.8 |
| 802 | Xanthoxyline | -5.9 |
| 803 | Xanthurenic Acid | -7.2 |
| 804 | XY 1 | -8.8 |
| 805 | Xylose | -5.5 |

| | | |
|---|---|---|
| 806 | Yohimbine HCl | -8.4 |
| 807 | Z JIB 04 | -8 |
| 808 | Zebularine | -6.3 |
| 809 | Zinc Undecylenate | -1.7 |
| 810 | β-thujaplicin | -6 |
